# miR-10a-5p as a critical molecular regulator of dopaminergic impulsivity in the nucleus accumbens

**DOI:** 10.64898/2026.07.20.739499

**Authors:** Yury Lages, Thibault Dufourd, Carole Carcenac, Mathilde Roux, Magali Bartolomucci, Frédérique Vossier, David Mallet, Robin Magnard, Colin Deransart, Sabrina Boulet, Pierre-Olivier Fernagut, Sebastien Carnicella

**Author notes:** YL and TD have equally contributed for this study. Corresponding author Sebastien Carnicella Grenoble Institut Neurosciences Univ. Grenoble Alpes Inserm, U1216 Chemin Fortuné Ferrini 38706 La Tronche Cedex, France.

## Abstract

Impulsive-compulsive disorders (ICDs), including pathological gambling, hypersexuality, and compulsive buying, are frequently precipitated by dopamine D2/3 receptor agonists such as pramipexole (PPX), yet the molecular mechanisms that confer individual vulnerability remain poorly understood. Impulsive choice, a core dimension of ICDs, is modulated by dopaminergic signaling within corticostriatal circuits, but the microRNAs (miRs) that potentially translate this signaling into persistent behavioral change have not been identified. Here, we combined a delay discounting task (DDT) with high-throughput miR sequencing in the dorsal striatum and nucleus accumbens (NAcc) of rats stratified by baseline impulsivity and subchronic PPX treatment. PPX increased impulsive choice selectively in low- and mid-impulsive rats, whereas high-impulsive rats remained unaffected, consistent with a ceiling effect. Among the differentially expressed miRs, miR-10a-5p emerged as the strongest candidate: it was constitutively elevated in high-impulsive rats and upregulated by PPX in low- and mid-impulsive animals in both regions, thereby paralleling the trait-dependent behavioral effect of the drug. In vivo viral-mediated overexpression of miR-10a-5p confirmed its predicted downregulation of the PI3K–AKT–mTOR and BDNF pathways in the striatum, and, critically, overexpression restricted to the NAcc, but not the dorsal striatum, was sufficient to increase impulsive choice, recapitulating the pro-impulsive effect of PPX. These findings identify miR-10a-5p as a critical molecular regulator of impulsivity through its activity in the NAcc, providing a mechanistic link between dopaminergic perturbation, trait vulnerability, and ICDs, and opening new avenues for the development of miR-directed therapeutic strategies for these disorders.

## Introduction

Impulsive-compulsive disorders (ICDs) encompass a spectrum of conditions characterized by a persistent impairment in the capacity to regulate reward-driven behaviors, leading to recurrent maladaptive patterns such as gambling, compulsive eating, hypersexuality, or excessive digital media use despite harmful consequences (1,2). The prevalence of ICDs, also conceptualized as behavioral addictions, in the general population can vary depending on the specific disorder, ranging from ∼1% in pathological gambling (3) to ∼5% in compulsive buying (4,5). Besides the direct individual and familial distress, ICDs are associated with increased rates of comorbid psychiatric conditions and reduced quality of life, highlighting substantial gaps in both detection and care. These gaps reflect our still incomplete understanding of the neurobiological mechanisms underlying ICDs; as a result, the few available treatment approaches are largely extrapolated from substance-use disorders and drug addiction models rather than grounded in ICD-specific pathophysiology (6,7).

Nonetheless, regardless of the specific behavioral expression of ICDs (gambling, hypersexuality, compulsive shopping, binge eating), impulsive choice, i.e. a systematic preference for small immediate over larger delayed rewards, appears to be a common psychological denominator of these disorders. This bias towards immediate gratification is tightly linked to dopaminergic modulation of mesocorticolimbic reward circuits, which contribute to encoding the value and timing of rewards and can bias choices toward immediate options (8,9). Consistently, iatrogenic ICDs frequently emerge as complications of dopamine replacement therapy in Parkinson’s disease (PD) and other dopamine-responsive conditions such as restless legs syndrome (RLS) and hyperprolactinemia (10,11). Specifically, D2/D3 receptor agonists such as pramipexole confer a markedly elevated risk of developing ICDs, with prevalence reaching 17.7% in PD patients (12), and ranging between 7.1% and 11.4% in RLS patients (13). Clinical studies and preclinical models demonstrate that pramipexole administration exacerbates impulsivity and maladaptive reward seeking, particularly by increasing delay intolerance in delay discounting tasks, a well-validated measure of cognitive impulsivity (14). However, only a subset of patients exposed to dopamine agonists develop ICDs, suggesting the existence of pre-existing vulnerability factors, such as baseline impulsivity or specific genetic and molecular profiles, conceivably similar to those implicated in drug addiction (9,15).

MicroRNAs (miRs), small non-coding RNAs that regulate protein-coding transcripts, represent one promising mechanism by which dopaminergic perturbations and individual vulnerability factors may translate into persistent maladaptive behaviors. miRs notably modulate gene networks that are central to synaptic remodeling and neuroadaptation (16,17), which are critically involved in reward processing and behavioral control (18,19). Consistent with this function, evidence from models of drug abuse shows that miRs modulate synaptic plasticity within corticostriatal circuits, particularly the nucleus accumbens (NAcc) and dorsal striatum (DS) (20,21), that may underlie impaired control over reward-seeking and persistent use despite negative consequences (19). These features being shared between substance use disorders and ICDs (16,22) within a common addiction framework (23,24) led us to hypothesize that dysregulation of specific miRs could contribute to the impulsive choice dimension of ICDs.

The present study therefore aimed to evaluate changes in miR profiles in a delay discounting task in rats following treatment with the dopamine agonist pramipexole. By stratifying animals according to baseline impulsivity and treatment condition, we identified miR-10a-5p in the DS and NAcc as consistently associated with high cognitive impulsivity, both as a trait and after pramipexole exposure. In addition, PI3K (phosphoinositide 3-kinase) and the downstream AKT/mTOR- or BDNF-related-pathways (protein kinase B, mammalian/mechanistic target of rapamycin, and brain-derived neurotrophic factor, respectively), which are known to modulate synaptic plasticity and to be implicated in drug-related addictive processes (25–27), were predicted as the most prominent targets of miR-10a-5p (28,29). We first confirmed in vivo, with a viral-mediated overexpression strategy, that miR-10a-5p indeed acts as a down-regulator of these pathways in the NAcc of rats. Finally, to establish a causal role, we showed that overexpression of miR-10a-5p in the NAcc was sufficient to increase impulsive behavior. Taken together, these findings identify miR-10a-5p as a critical molecular regulator of cognitive impulsivity. They further suggest that miR-dependent modulation of NAcc signaling may provide a mechanistic link between dopaminergic perturbations, trait impulsivity, and vulnerability to ICDs and related behavioral addictions.

## Methods

### Animals

Male Sprague Dawley rats (Janvier, France) were used for all experiments. For behavioral experiments (Experiments 1 and 2), animals were 8 weeks old at the start of the protocol, were euthanized no later than 18 weeks of age, and were maintained under a restricted feeding regimen (85-90% of standard body weight) throughout testing. For molecular pathway analysis (Experiment 3), rats were 12 weeks old at the time of surgery with no food restriction. The exact number of rats included in each experimental protocol is specified in the corresponding methods subsection. All procedures followed European Union and French ethical guidelines for animal research (Directive 2010/63); further details on housing conditions and ethical approval are provided in the Supplementary Methods.

### Delay discounting procedure

Impulsive choice was assessed using a delay discounting task (DDT) adapted from Magnard et al (30). Experiments were conducted in standard operant conditioning chambers (Med-Associates) equipped with two retractable levers, which delivered 5% or 10% (w/v) sucrose solutions upon lever press. Training and testing sessions were divided into 3 experimental phases and are described in detail in the Supplementary Methods.

After the three phases of the DDT, the animals were separated based on a quartile approach, as previously used for the evaluation of impulsive traits (31). Specifically, they were divided into three groups according to their preference for the larger reward over the smaller one at the end of the last delay discounting session: low-impulsive (LI; ≥ 75% preference for the large reward), mid-impulsive (MI; 25% ≤ preference < 75%) and high-impulsive (HI; ≤ 25% preference for the large reward). They were then subjected to a subchronic treatment with pramipexole or viral infection for miRNA overexpression. Subsequently, the animals were reevaluated in the DDT starting at phase 2. Impulsive choice was indexed by the area under the curve (AUC) of preference for the large reward across delays and by the hyperbolic discounting rate k (32); calculation details are provided in the Supplementary Methods.

### Experimental design

This study was structured into three main experiments, which are briefly described below and with more detains in the Supplementary Methods.

#### Experiment 1: Subchronic treatment with pramipexole

Forty-eight male rats selected after DDT received daily intraperitoneal injections of either pramipexole or vehicle during the second evaluation of DDT (Figure 1); brains were collected 24 h after the last DDT session.

**Figure 1.**
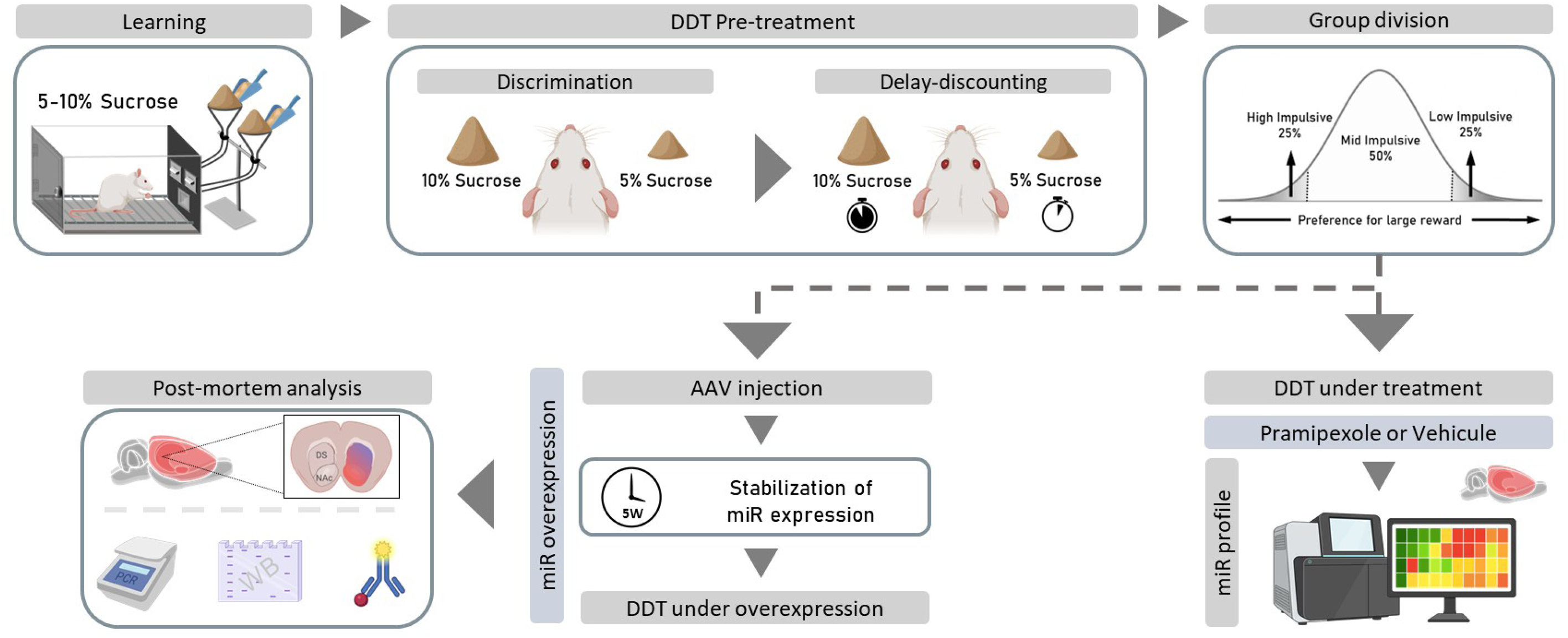

#### Experiment 2: Overexpression of microRNA constructs

AAV vectors encoding miR-10a-5p or a negative control miR were bilaterally injected into the NAcc core of 119 naïve rats; subsets of brains were collected at 2, 5, or 8 weeks post-surgery to assess expression kinetics, and at 5 weeks to assess miR-10a-5p target expression.

#### Experiment 3: Causal implication of miR-10a-5p in impulsive behaviors

A separate cohort of 104 rats, phenotyped by the DDT and stratified by impulsivity level, received the same viral constructs in the NAcc or DS and was retested in a second round of the DDT after 5 weeks of viral expression; brains were collected 24 h after the last DDT session (Figure 1) and processed as described in the Supplementary Methods.

### microRNA extraction

Total and small RNA (<200 nucleotides) were isolated from brain tissue (up to 50 mg) using phenol-chloroform-based commercial kits (MiRNeasy Mini Kit and MiRNeasy MinElute Cleanup Kit, Qiagen). Biopsies from both brain hemispheres were pooled to ensure a sufficient amount of starting material.

### High-throughput sequencing, RT-qPCR, and Western blot analysis

Small RNA-enriched brain samples were sequenced (QIAseq miRNA Library Kit, NextSeq 500) to profile miR-10a-5p expression across impulsivity subgroups, and results were validated by RT-qPCR. mRNA expression of PI3K and BDNF was also assessed by RT-qPCR. Protein levels of AKT, p-AKT, RPS6, and p-RPS6 were quantified by Western blot. Full protocols are provided in the Supplementary Methods.

### Gene ontology

Gene Ontology (GO) and pathway enrichment analysis of the predicted targets of miR-10a-5p was performed using the g:GOSt function of the g:Profiler online platform (https://biit.cs.ut.ee/gprofiler/gost). For more details, refer to the Supplementary Methods. GO terms were queried within the Biological Process (BP) and Molecular Function (MF) namespaces, and pathway enrichment was evaluated using Kyoto Encyclopedia of Genes and Genomes (KEGG) annotations.

### Statistical analysis

#### Sample size determination

Sample sizes were determined separately for each experimental component. For the behavioral cohort and the assessment of miR-10a-5p effects on impulsive choice, sample size was determined a priori using G*Power* (version 3.1) for a two-way repeated-measures ANOVA (delay × viral vector; within–between interaction). The effect size was estimated from the pramipexole-induced change in delay discounting AUC in a pilot experiment and cross-validated against Magnard et al. (30). Other fixed parameters were: r = 0.5, ε = 0.75, α = 0.05, power = 0.8-0.9. A first experimental cohort further confirmed that the target n per subgroup was achievable given the expected proportion of high- and low-impulsive animals. For the miR-10a-5p kinetics and RT-qPCR/western blot experiments, group sizes were additionally constrained by histologically confirmed viral injection placement and adequate tissue yield. For the high-throughput sequencing experiment, n = 6 per subgroup was used given the multi-subgroup design (three impulsivity levels × two treatment conditions, in two brain regions). This reduced n was compensated by an analytical pipeline that selected candidate miRNAs based on expression amplitude (VIP > 1.5) and within-group homogeneity, thereby restricting detection to the most robust signals. This strategy was not designed to detect all differentially expressed miRNAs, but to identify the most robust ones — a trade-off discussed further in the Discussion.

#### Behavioral assessments

All results are presented as mean ± SEM. Statistical analyses were performed using SigmaStat (version 4.0, Systat Software Inc., USA) and GraphPad Prism (version 9.0.2, USA), with a significance threshold of α = 0.05. Delay discounting performance was analyzed using two-way repeated-measures ANOVAs, with delay as the within-subject factor and impulsivity group and/or treatment as between-subject factors. Geisser–Greenhouse corrections were applied when the sphericity assumption was violated, and significant main effects and interactions were followed up using Newman–Keuls post-hoc tests.

#### Molecular analyses

For high-throughput sequencing, quality control was performed by Genosplice (ICM, Paris). Data normalized via edgeR in R (v.3.2.5) were processed using SIMCA and Microsoft Excel. A global multivariate analysis was first conducted using unsupervised principal component analysis (PCA) on the normalized miRNA expression matrix to explore overall patterns of variance across treatment and impulsivity levels. Unsupervised PCA identified microRNAs driving separation between groups defined by treatment (pramipexole or vehicle), baseline impulsivity level (high, mid, low), or both. MicroRNAs with a variable importance in projection score > 1.5 were retained as predictive, and the resulting lists were cross-referenced to identify miRNAs robustly associated with each condition. These candidates were then tested in GraphPad Prism (version 9.0.2, USA) with two-way ANOVA (factors: treatment × impulsivity level) with Bonferroni correction; post-hoc pairwise comparisons used Bonferroni-corrected t-tests. The same statistical approach was applied to RT-qPCR data for miR-10a-5p kinetics. For RT-qPCR quantification of *Pi3k* and *Bdnf* mRNA levels and for western blot protein quantification, comparisons were restricted to the miR-10a-5p and miR-Neg groups using two-tailed unpaired Student’s t-tests. Effect sizes are reported as partial eta-squared (η²p), calculated as η²p = SS_effect / (SS_effect + SS_residual) for ANOVA-based analyses and as η²p = t²/(t² + df) for t-test comparisons. Effect size magnitude was interpreted as follows: η²p < 0.06 indicates a small effect, 0.06 ≤ η²p < 0.14 indicates a medium effect, and η²p ≥ 0.14 indicates a large effect (33,34). Because RT-qPCR measurements were obtained across cohorts run at different time points with different reagent batches, expression values were z-score normalized before analysis to control for inter-batch variance; otherwise, all results are presented as mean ± SEM.

## Results

### Pramipexole-induced impulsivity depends on baseline trait

Following the delay discounting task, different profiles of preference for the large reward were observed [interaction delay × trait: F6_,135_= 18.4, p < 0.0001; effect of trait: F_2,45_ = 114, p < 0.0001] (Figure 2A). The animals were then classified according to baseline cognitive impulsivity into three distinct groups: high- (HI), mid- (MI), and low-impulsive (LI), using a quartile-based approach based on the analysis of mean AUC values [F_2,45_= 25.93, p < 0.0001] and confirmed by the discounting rate k [F_2,45_= 11.17, p < 0.0001] (Figure 2B-C). Subgroups were formed and counterbalanced for subsequent pramipexole or vehicle treatment, with no pre-existing differences between treatment conditions [AUC: effect of trait between groups: F_2,42_= 26.71, p < 0.0001; effects of treatment or treatment × delay interaction: Fs < 2.12, ps > 0.133 between subgroups].

**Figure 2.**
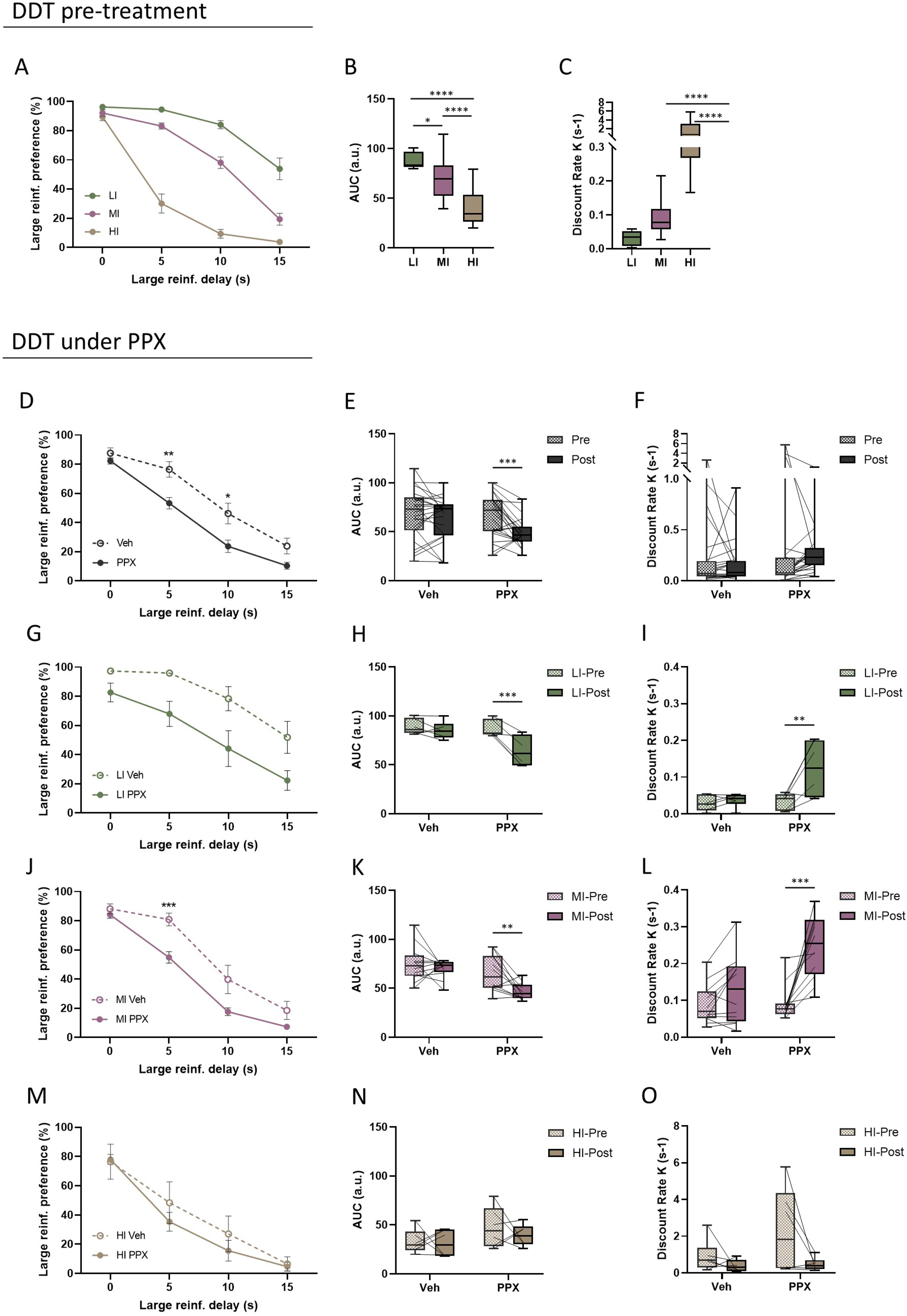

Subchronic pramipexole treatment increased cognitive impulsivity relative to vehicle, as reflected by a greater decline in preference for the large reward with the increase in delay [effect of treatment: F_1,46_ = 9.28, p = 0.004; treatment x delay interaction: F_3,138_ = 3.91, p = 0.010] (Figure 2D), and confirmed by AUC [interaction delay × treatment: F_1,46_ = 5.74, p = 0.021] (Figure 2E) and discounting rate k (Figure 2F) [effect of trait: F_2,42_ = 3.80, p < 0.0001]. Critically, this effect was trait-dependent: pramipexole increased impulsivity selectively in LI and MI animals [treatment x delay interaction of MI: F_3,66_ = 3.90, p = 0.013; effect of treatment: F_s_ > 8.49, ps < 0.013], as confirmed by both AUC [interaction delay x treatment: F_s_ > 10.48, ps < 0.009; effect of treatment: MI: F_1,22_ = 8.49, p = 0.008] and discount rate k [interaction delay x treatment: LI: F_1,10_ = 7.75, p = 0.019; effect of treatment: MI: F_1,22_ = 11.64, p = 0.003], but had no significant effect in HI animals [preference for large reward, treatment x delay interaction: F_3,30_ = 0.543, p = 0.657; effect of treatment: F_1,10_ = 0.429, ps = 0.5272] (Figure 2G-O). Taken together, these findings indicate that subchronic pramipexole exacerbates delay intolerance preferentially in rats with low and intermediate baseline impulsivity, revealing a trait-dependent effect of the drug on cognitive impulsivity.

### miR-10a-5p links pramipexole to impulsivity

To identify microRNAs associated with pramipexole treatment, impulsivity trait, or their interaction, high-throughput sequencing was performed on DS and NAcc tissue collected from animals used in Experiment 1 (see Methods). All analyses were performed without prior assumptions to prevent target selection bias. Raw sequencing output was processed for quality control, adapter removal, and alignment to the rat genome (848 known sequences, Figure 3A). Read counts corresponding to each microRNA were quantified for each sample, and expression was normalized per library (reads per million). To select biologically relevant microRNAs (highest robustness), a minimum expression-level threshold was established: microRNAs were retained if normalized expression reached 15 reads per million in at least one experimental condition (standard normalization typically detects microRNAs at below 1 read). Only microRNAs expressed in at least 50% of an experimental condition (≥3 of 6 animals) were retained for further analysis. This yielded 337 microRNAs in the DS, and 349 in the NAcc. Among these microRNAs readily expressed in the striatum, a global multivariate analysis (see Methods) identified 203 targets in the DS, and 209 in the NAcc that contribute to the discrimination of at least the impulsive trait and/or the treatment as experimental conditions. 29 microRNAs in the DS, and 8 in the NAcc of these selected targets were further statistically confirmed with two-way ANOVA analyses (Figure 3A). Interestingly, treatment effects were predominantly found in the DS, whereas trait-related effects were more evident in the NAcc (Figure 3B). Among these validated targets, all were conserved in humans except miR-672-3p, miR-673-5p, miR-1188-5p, and miR-3068-5p, which were therefore excluded for further analyses.

**Figure 3.**
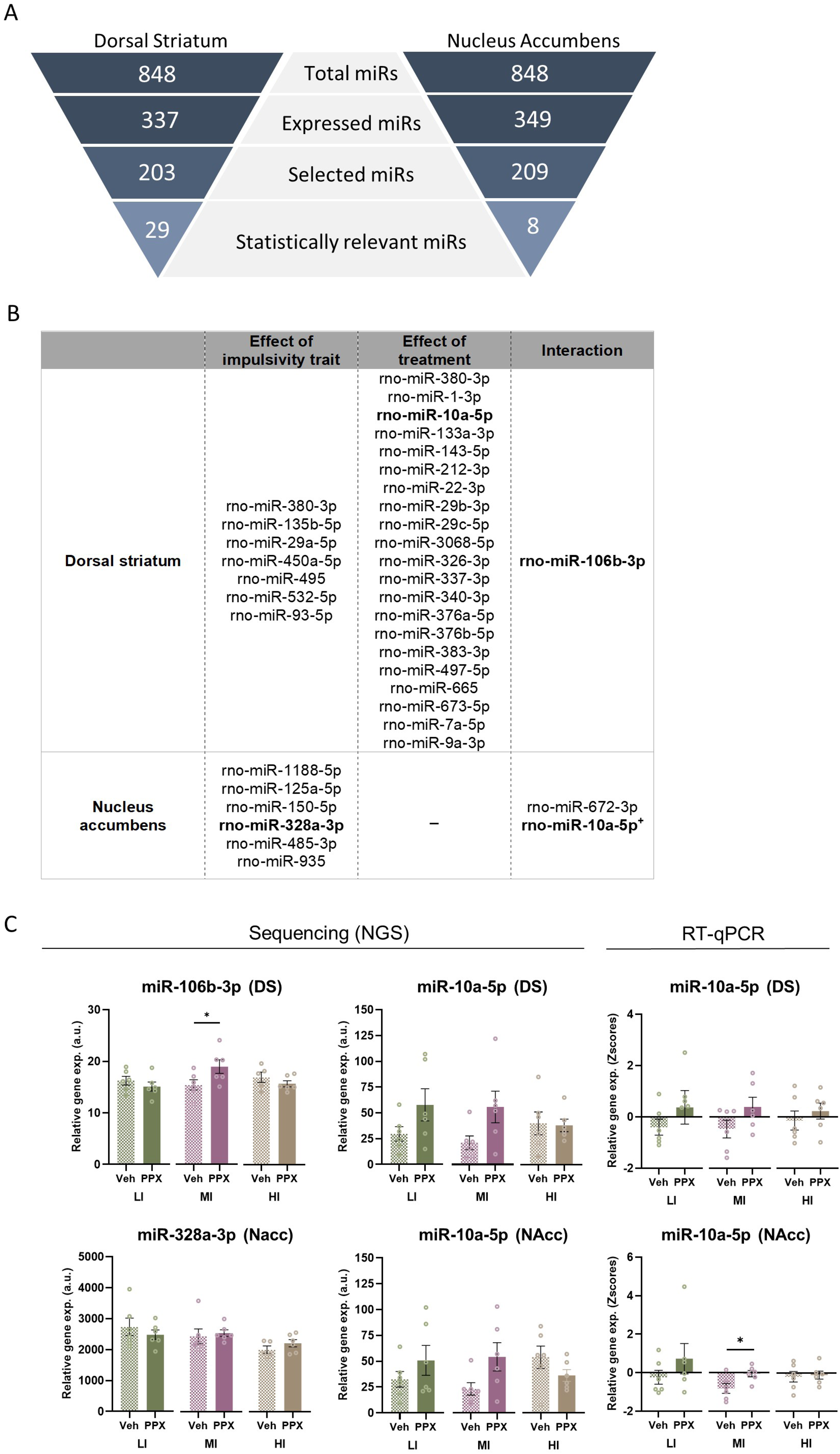

The filtering pipeline was completed with post-hoc testing to further validate the localized dysregulations, with representative examples shown in Figure 3C. miR-106b-3p in the DS showed, for instance, a significant trait × treatment interaction [F_2,29_ = 4.09, p = 0.027, η²p = 0.220], reflecting a selective increase following pramipexole specifically in MI animals [t(29) = 2.63, adjusted p = 0.04, η²p = 0.193], with no significant modulation detected in the NAcc. However, as pramipexole increased impulsivity in both LI and MI animals, the selective upregulation of miR-106b-3p in MI animals only partially mirrored the behavioral data, and this miRNA was therefore not retained for further investigation. Similarly, miR-328a-3p in the NAcc showed differential expression across impulsivity levels [F_2,29_ = 3.84, p=0.033, η²p = 0.209], independent of treatment [F_1,29_ = 0.018, p = 0.895, η²p < 0.001], with a trend for higher expression in LI than HI rats (Bonferroni-corrected trend, adjusted p = 0.07), suggesting a putative role as a marker of baseline impulsivity trait rather than drug responsiveness, and was equally set aside.

In contrast, miR-10a-5p demonstrated a particularly relevant profile. Sequencing from the DS revealed a significant main effect of pramipexole treatment on its expression [F_1,30_ = 5.10, p = 0.031, η²p = 0.096], with a directional pattern in which HI animals showed persistently elevated levels regardless of treatment, while LI and MI animals showed higher expression following pramipexole administration (Figure 3B). Although the trait × treatment interaction did not reach statistical significance [F_2,30_ = 1.59, p = 0.222, η²p = 0.145], likely reflecting limited statistical power, this pattern closely mirrored the trait-dependent behavioral effect of pramipexole. A stronger trend toward a trait × treatment interaction was also observed in the NAcc [F_2,30_ = 2.98, p = 0.066, η²p = 0.166]. These sequencing findings were then confirmed by RT-qPCR, which yielded a significant treatment effect in the NAcc [F_1,30_ = 6.12, p = 0.019, η²p = 0.169] and a trend in the DS [F_1,30_ = 3.92, p = 0.057, η²p = 0.115], without significant trait × treatment interactions [F_s_ < 0.76, p > 0.475, η²p < 0.048]. The consistency of this expression profile across both techniques and both brain regions, and its alignment with the trait-dependent behavioral effects of pramipexole, thereby led us to prioritize miR-10a-5p for causal investigation.

### miR-10a-5p targets addiction-related signaling pathways

We next investigated the predicted molecular targets of miR-10a-5p and their biological relevance through bioinformatic analysis. Fifty-three possible targets of this miRNA were identified (Supplementary Figure 3). Gene Ontology enrichment analysis indicated that most targeted genes participate in the positive regulation of cellular and biological processes, including primary, macromolecule and RNA metabolic pathways (Figure 4A; BP1–BP7). In addition, significant enrichment was observed for Molecular Function terms related to enzyme and protein binding (MF1–MF2), and for a KEGG pathway associated with proteoglycans in cancer, supporting the involvement of miR-10a-5p targets in core signaling and oncogenic processes (Figure 4A).

**Figure 4.**
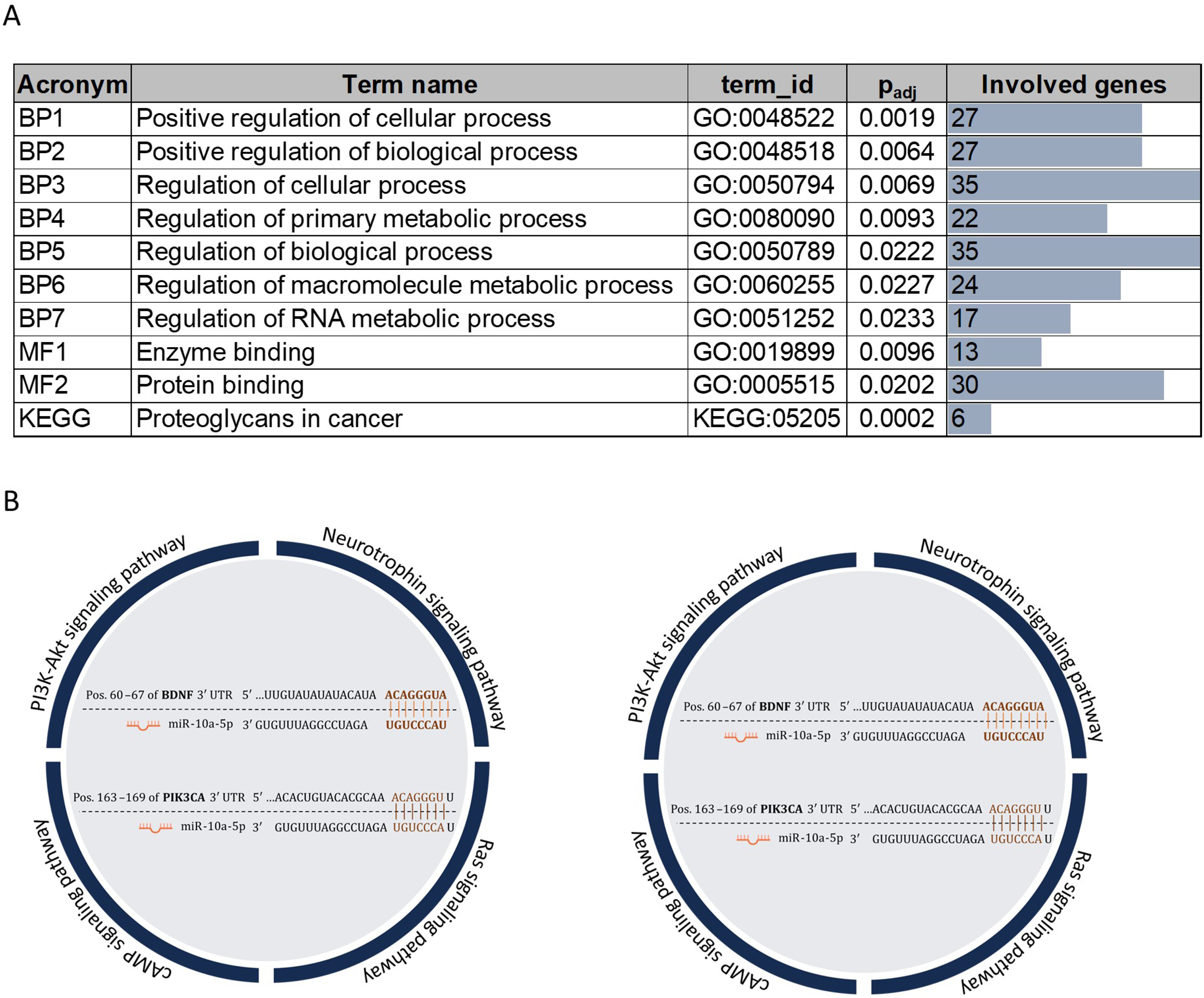

Among its predicted targets, *Pi3k* (35,36) and *Bdnf* (37) have already been validated in cancer-related studies and miR-10a-5p has also been linked to Bdnf regulation in prefrontal cortical and hippocampal circuits (38,39). miR-10a-5p binds to *Pi3k* mRNA at positions 163–169 bp, and to *Bdnf* mRNA at positions 65–72 bp, leading to their degradation (Figure 4B) and thereby reducing the expression of these genes. Moreover, downregulation of these targets by miR-10a-5p is further predicted to suppress mTORC1 activity (36,40).

### miR-10a-5p modulates BDNF and PI3K-related signaling cascades

Because these candidates, and their related signaling axis, have been previously shown to be strongly implicated in drug-induced neuroadaptations (41,42), we next aimed at confirming that miR-10a-5p acts indeed as a downregulator of these pathways *in vivo*, with a viral-mediated overexpression of this miR in the core of the NAcc (Figure 5A), as this subregion is known to be central in the regulation of cognitive impulsivity (43,44).

**Figure 5.**
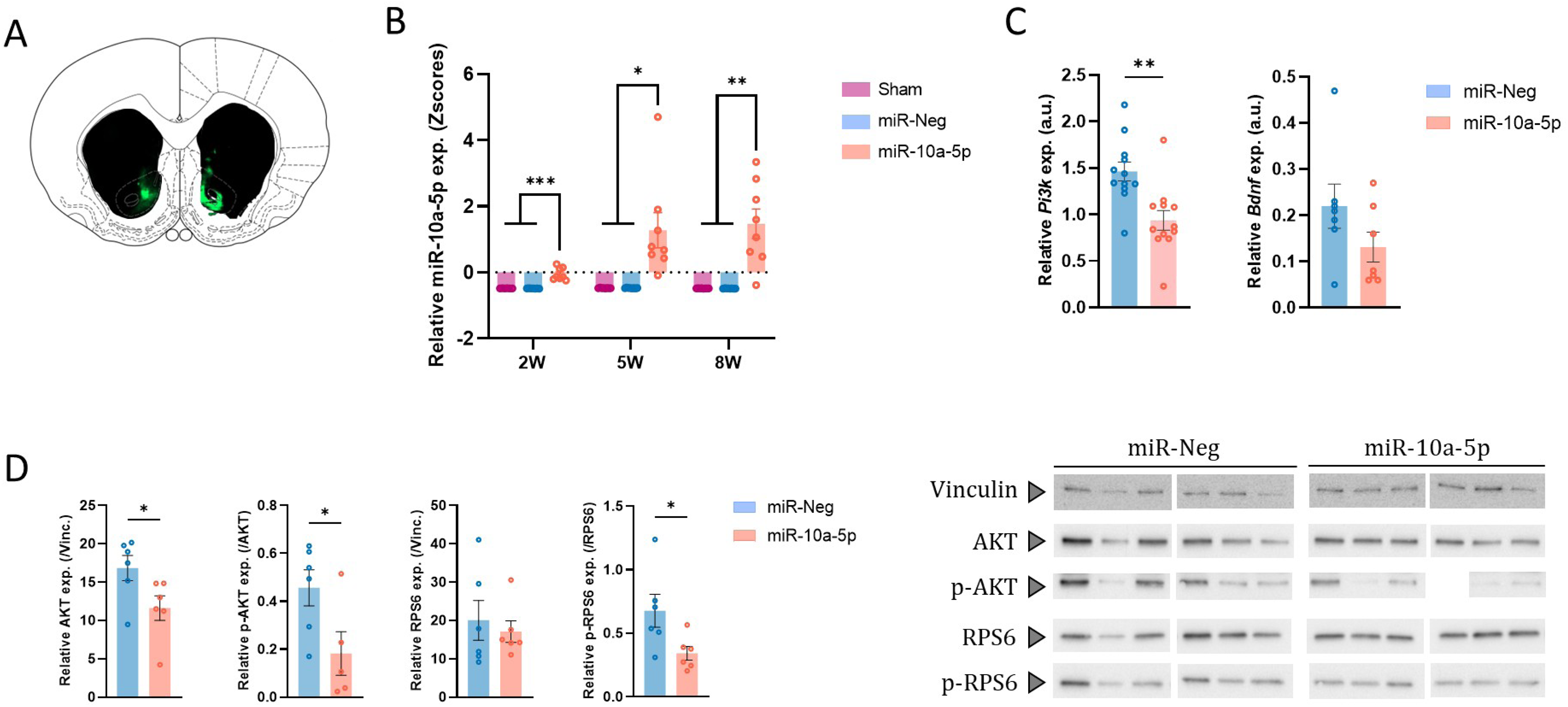

Following the confirmation of the overexpression of miR-10a-5p viral construct and analysis of its time-course (Figure 5B), we then tested whether miR-10a-5p regulates two of its *in silico* predicted targets, *Pi3k* and *Bdnf*, in the NAcc. RT-qPCR confirmed that miR-10a-5p overexpression led to a significant decrease in the expression of *Pi3k* mRNA [t(22) = 3.59, p = 0.002, η²p = 0.369] and to a lesser extent of *Bdnf* mRNA [t(12) = 1.54, p = 0.150, η²p = 0.165] in this structure (Figure 5C). We then investigated whether the downregulation in the expression of *Pi3k* and *Bdnf* would effectively impact shared downstream pathways, such as AKT and RPS6 through AKT-mTORC1 axis at protein level (see discussion) (45,46). Quantification by western blot indicated that, in the miR-10a-5p group, AKT and its active form (p-AKT) had decreased expression [AKT: t(10) = 2.28, p = 0.046, η²p = 0.342; p-AKT: t(9) = 2.35, p = 0.044, η²p = 0.379] (Figure 5D). Also, there was a decreased phosphorylation of RPS6 [t(10) = 2.38, p = 0.039, η²p = 0.315], while the inactive form remained unchanged [t(10) = 0.5, p = 0.630, η²p = 0.024] (Figure 5D). Taken together, these results demonstrate the biological significance of miR-10a-5p as a negative modulator of this canonical pathway associated with addictions. They thereby further support the relevance of miR-10a-5p as a candidate to investigate, and validate the developed overexpression strategy *in vivo*.

### miR-10a-5p overexpression in NAcc drives impulsivity

To test whether miR-10a-5p plays a causal role in impulsive behavior, we next overexpressed it in the NAcc of rats performing the delay discounting task. Similarly to the behavioral protocol applied to test the effects of PPX, naïve animals went through a first round of DDT, at the end of which they were segregated according to their levels of impulsivity. LI and MI animals, then, received either miR-Neg or miR-10a-5p AAV injections in the NAcc. Post-mortem analyses confirmed that miR-10a-5p overexpression targeted mainly the core region of the NAcc (Figure 6A), leading to significant increased expression in miR-10a-5p AAV-injected rats (Figure 6B). 5 weeks after surgeries, rats were retested in the DDT. Post-infection behavioral testing showed a general reduction in the preference for the larger reward in the miR-10a-5p group [miR x delay interaction: F_3,141_ = 2.766, p = 0.044; effect of miR: F_1,47_ = 3.051, p = 0.087] (Figure 6C). In low-impulsive animals, this effect did not translate into a clear change in the preference for the larger reward across delays [F_s_ < 0.429, ps > 0.181] (Figures 6D-E), although miR-10a-5p overexpression significantly increased the individual discount rate k after infection [Pre x Post, miR-10a-5p group, t(17) = 2.69, adjusted-p = 0.03] (Figure 6F), indicating that LI rats infused with miR-10a-5p discounted delayed rewards more steeply than miR-Neg controls. By contrast, in MI animals, miR-10a-5p overexpression induced a marked increase in delay discounting, as evidenced by a stronger decline in preference for the larger reward across delays compared with the miR-Neg group [interaction miR x Delay: F_3,72_ = 4.78, p = 0.004; effect of miR: F_1,24_ = 3.664, p = 0.068] (Figure 6G). This difference was maintained when we compared the AUC [interaction miR x Time: F_1,24_ = 7.36, p = 0.012] of pre- and post-infection DDT profiles of MI rats (Figure 6H), although the discount rate k did not change significantly in this subgroup [interaction miR x Time: F_1,24_ = 1.90, p = 0.181] (Figure 6I).

**Figure 6.**
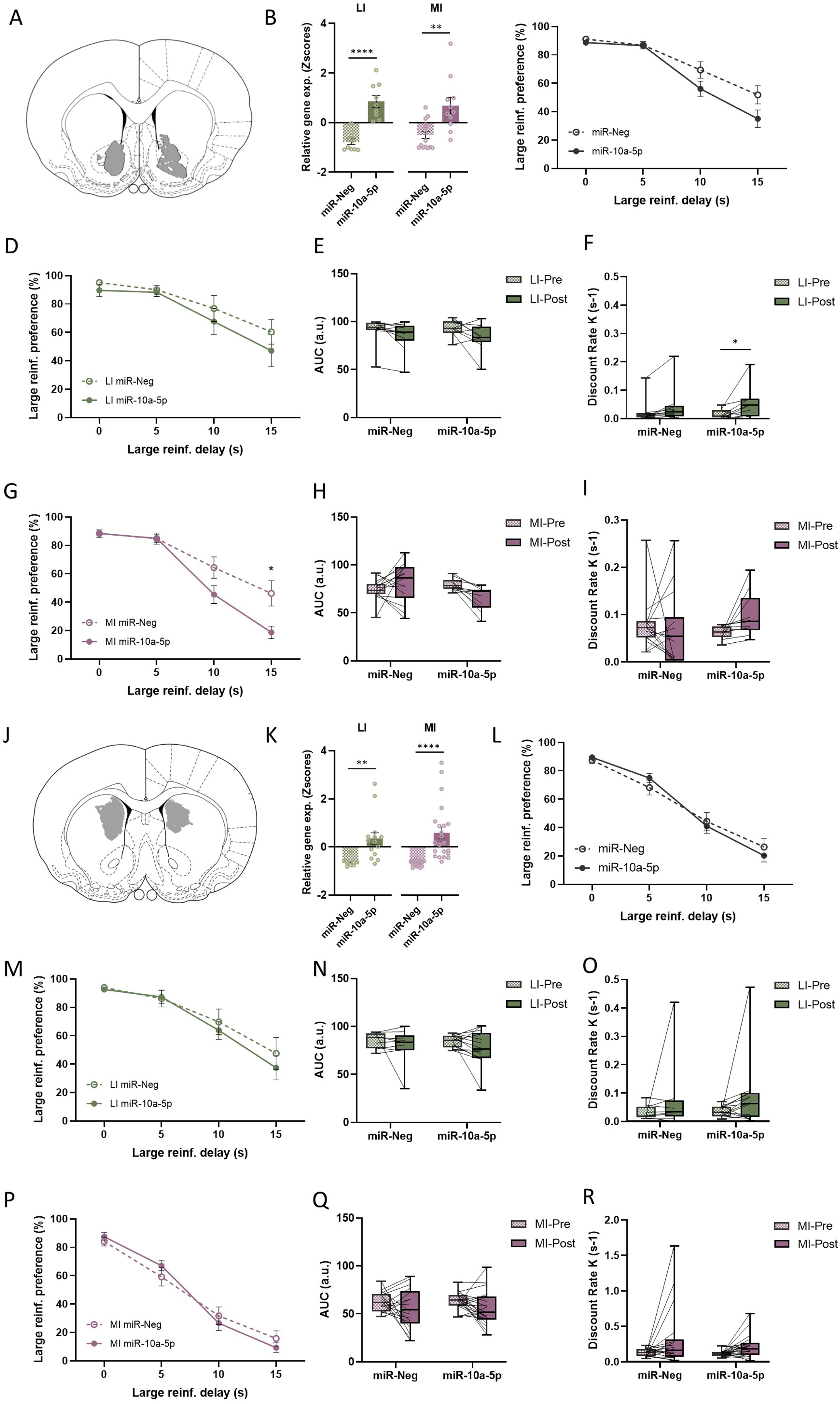

Because the dorsomedial striatum (DMS) has also been implicated in cognitive impulsivity (47,48) and pramipexole-induced upregulation of miR-10a-5p was also observed in the DS, we next examined whether overexpression of miR-10a-5p in the DMS (Figure 6J) influences impulsive behavior. However, even though an increased expression of miR-10a-5p was verified in the DMS at the end of the experiment (Figure 6K), no changes in behavior were observed in either low- or mid-impulsive rats [Fs < 1.637, ps > 0.208] (Figure 6L-R). Altogether, the evidence supports a key involvement of miR-10a-5p in cognitive impulsivity in rats, primarily via its activity in the NAcc.

## Discussion

In the present study, trait-dependent vulnerability and pramipexole exposure were associated with broad microRNA alterations in both DS and NAcc, with distinct profiles related to baseline impulsivity, pharmacological treatment, or their interaction. Among these candidates, miR-10a-5p emerged as a potential marker of high impulsivity. Indeed, it appears constitutively elevated in high-impulsive rats and upregulated by pramipexole in low- and mid-impulsive animals in both striatal regions, in parallel with the drug-induced increase in impulsive choice observed specifically in the least impulsive (LI and MI) rats. Viral-mediated overexpression of miR-10a-5p in the NAcc core principally, but not in the DS, was sufficient to increase impulsive choice, thereby recapitulating the pramipexole-induced behavioral phenotype. At the molecular level, miR-10a-5p overexpression decreased *Pi3k* and *Bdnf* mRNA expression and reduced AKT and mTORC1 signaling in the NAcc, as indicated by lower phosphorylation of RPS6, confirming the functionality of the construct and pointing to a canonical AKT–mTOR pathway as a downstream effector previously implicated in drug-induced neuroadaptations and maladaptive reward seeking. Together, these results support a model in which miR-10a-5p acts as a convergent node linking dopamine agonist exposure, striatal circuit specificity, and cognitive impulsivity relevant to ICDs.

This identification of miR-10a-5p as a candidate regulator of dopaminergic impulsivity was the result of a stepwise microRNA profiling approach. We first performed small RNA sequencing in the DS and NAcc of rats stratified by baseline impulsivity and pramipexole treatment. Given the multi-subgroup design, sequencing was performed with a reduced n per subgroup, which may have limited sensitivity to detect subtle effects but favored the detection of robust, consistent signals, an important prerequisite to move from correlational to causal investigations. After quality control, expression filtering and multivariate analyses, a restricted set of dysregulated miRNAs was retained and further evaluated by statistical analysis and RT-qPCR. Some candidates showed associations with trait or treatment, but their expression patterns did not consistently match the pramipexole-induced changes in impulsive choice and were not considered for further experiments. In contrast, miR-10a-5p showed a coherent profile across regions and analyses, with evidence that its expression is associated with increased impulsivity, whether driven by an innate trait or by pramipexole treatment, making it the principal candidate for causal investigation.

In the delay discounting task, high-impulsive rats did not show a further pramipexole-induced increase in impulsive choice, which most likely reflects a ceiling effect, as these animals already displayed near-maximal preference for immediate rewards at intermediate delays. This observation is in line with the miR-10a-5p expression profile: levels are higher in high-impulsive rats than in less impulsive animals at baseline, while pramipexole tends to increase miR-10a-5p expression in low- and mid-impulsive animals in both DS and NAcc. Pramipexole therefore shifts low- and mid-impulsive rats toward a more “high-impulsive-like” miR-10a-5p expression pattern, particularly in the NAcc, suggesting a partial molecular convergence between trait- and treatment-related vulnerability that may help explain why dopamine agonists can precipitate ICDs in only a subset of patients, including also those without prominent baseline impulsive traits (13,49).

Beyond impulsivity, miR-10a-5p has been previously implicated in the regulation of Bdnf and its downstream signaling targets in the hippocampus and prefrontal cortex, in contexts such as peripubertal alcohol exposure and mood-related phenotypes (38,39,50). Two of its predicted targets, Pi3k and Bdnf, have also been experimentally validated as direct miR-10a-5p targets in cancer models (28,37). These findings converge with broader evidence that BDNF–TrkB and PI3K–AKT–mTOR cascades regulate synaptic plasticity and are critically involved in addiction-related neuroadaptations (26,27,38,40,51). BDNF–TrkB and PI3K converge onto the same PI3K–AKT–mTORC1 axis, such that BDNF binding to TrkB promotes recruitment of PI3K and activation of AKT, which in turn drives mTORC1-dependent phosphorylation of RPS6 as a key translational effector (52,53). Consistent with these data, overexpression of miR-10a-5p in the NAcc in our model decreased *Pi3k* and *Bdnf* mRNA expression and reduced AKT and mTORC1 signaling, as indicated by lower levels of AKT and p-AKT and reduced phosphorylation of RPS6, while total RPS6 remained unchanged. The concomitant reduction in AKT protein and phosphorylation suggests that miR-10a-5p not only dampens pathway activity via *Pi3k* downregulation but may also decrease AKT expression itself, either through direct regulation or indirect modulation within the PI3K–AKT cascade. Together, these findings demonstrate that miR-10a-5p can gate a canonical neurotrophic-related signaling pathway in a striatal subregion directly involved in dopaminergic impulsivity and addiction-relevant decision processes. Given the established role of AKT–mTOR signaling in drug-induced neuroadaptations and maladaptive reward seeking, such modulation provides a plausible mechanism contributing to the observed increase in delay intolerance. However, the present experiments were not designed to specifically determine whether AKT–mTOR- and BDNF-related changes are necessary or sufficient for the behavioral effects of miR-10a-5p, and other targets of this microRNA may also contribute to its impact on impulsive choice.

In line with this mechanistic role in NAcc circuits, experimentally induced overexpression of miR-10a-5p in the NAcc increased cognitive impulsivity, as shown by a steeper decline in preference for the larger delayed reward and a reduction in AUC in mid-impulsive rats, and by an increased discount rate k in low-impulsive rats. These effects closely resembled the pramipexole-induced increase in delay intolerance observed in low- and mid-impulsive animals, indicating that elevated miR-10a-5p levels in the NAcc are sufficient to reproduce the main behavioral features of the dopaminergic D3/D2 stimulation, but with some differences depending on impulsive basal levels. In contrast, although miR-10a-5p overexpression in the dorsomedial striatum produced a robust increase in miR-10a-5p levels, it did not alter performance in the DDT in either low- or mid-impulsive rats. This dissociation supports a model in which miR-10a-5p exerts a causal influence on impulsive choice primarily through its activity in NAcc-associated circuits.

The regional specificity of the behavioral effects of miR-10a-5p overexpression is consistent with the distinct functional roles of NAcc and DS in adaptive behavior. The NAcc, and particularly its Core subdivision, mainly targeted here, is critically involved in reward valuation, temporal discounting, and the integration of motivational signals that direct choices toward immediate or delayed options, processes directly engaged by the DDT (44,54). The DS, in contrast, is functionally divided into the DMS and the dorsolateral striatum (DLS), which contribute to distinct aspects of behavioral control. While the DMS has also been implicated in impulsivity through its role in goal-directed learning and action-outcome associations, it operates together with the DLS in the control of adaptive behavior, and dysfunctions in this system are more closely associated with the emergence of compulsive patterns of responding, (55,56). Accordingly, the absence of a behavioral effect of DS miR-10a-5p overexpression in the DDT does not rule out a contribution of this structure to other aspects of ICDs more related to its compulsive dimension, such as perseverative responding, which would require complementary behavioral paradigms to be revealed.

Elucidating microRNA function in ICDs is expected to advance the still very limited understanding of the molecular mechanisms underlying behavioral addictions in humans. By dissecting how specific regulatory miRs, such as miR-10a-5p, may act as a molecular gate to reshape corticostriatal circuits to promote impulsive choice, the present work provides causal evidence that microRNA dysregulation can control an addiction-relevant behavioral dimension in a non-substance domain, thereby extending to ICDs molecular principles previously established in drug addiction models. This mechanistic insight refines the pathophysiological framework of ICDs, in which dopamine agonist exposure interacts with molecular vulnerability factors within NAcc and dorsal striatal networks, and delineates new trajectories for therapeutic development at a time when miR-directed strategies such as antagomiRs and miRNA mimics are currently being developed in other medical fields (57,58). In this context, future studies should investigate the potential of miR-10a-5p as a circulating biomarker of ICD risk in dopamine agonist-treated patients and possibly in other behavioral addictions, and should assess whether targeting miR-10a-5p might be a valuable therapeutic strategy for the treatment of impulsivity- and compulsivity-related disorders.

## Acknowledgements

We thank the GIN behavioral facility, which is supported by the Grenoble Center of Excellence in Neurodegeneration (GREEN), Julie Brocard for her help with cell culture and the PIC GIN microscopy imaging platform.

## Author contributions

YL, TD, and SC designed the experiments with CD, SB and POF. YL and TD performed experiments and analyses with CC, MR, MB, FV, DM and RM. YL, TD, and SC wrote the manuscript with inpu from the other coauthors.

## Funding

This work was supported by the Institut National de la Santé et de la Recherche Médicale (INSERM), Association France Parkinson (to TD and SC), Agence Nationale de la Recherche (ANR) (ANR13 SAMA001401 and ANR-21-CE37-0013 DISCOMMODE to SC, and ANR-22-CE17-0011 DOPAMINE to POF), Institut National du Cancer and Institut de Recherche en Santé Publique (INCA-IReSP miRACle to SC and POF), Fondation de France (to SC), fundings managed by the National Research Agency under France 2030 (ANR-22-EXPR-0010 to YL), and Grenoble Alpes University (particularly the Pôle Universitaire d’Innovation – PUI).

## Conflict-of-interest statement

We declare no conflict of interest.

---

**Supplementary Figure 1.**
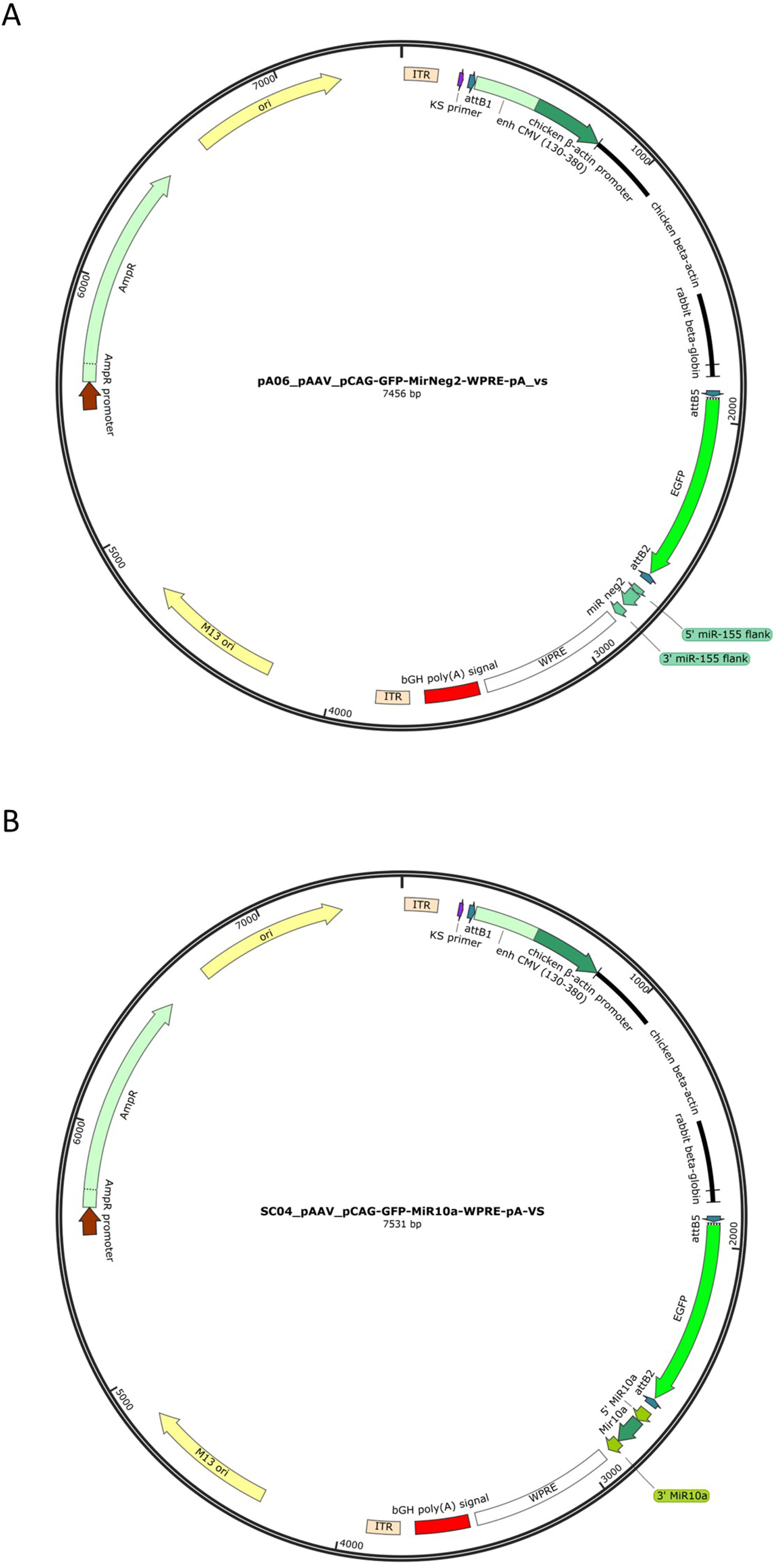

**Supplementary Figure 2.**
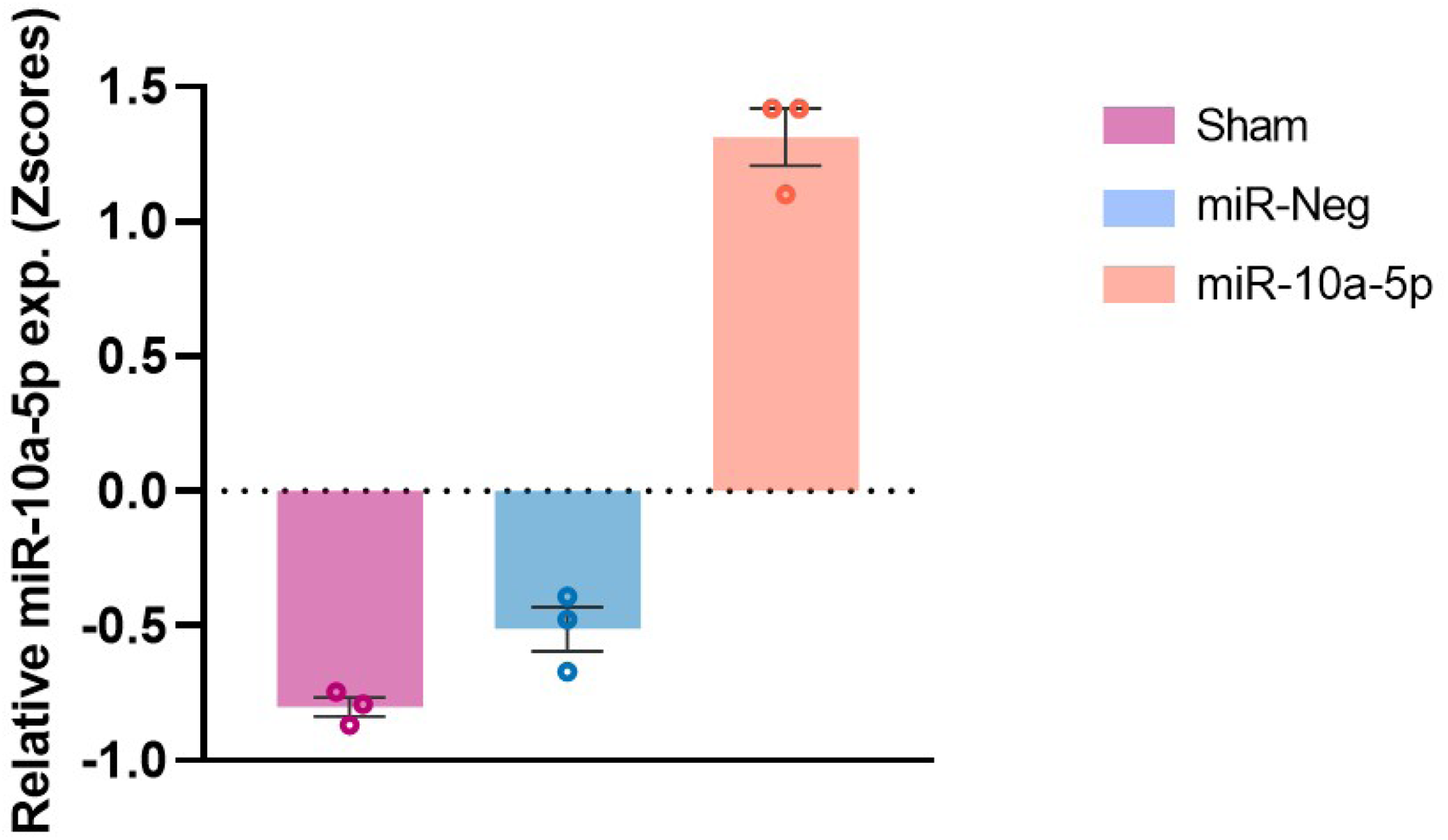

**Supplementary Figure 3.**
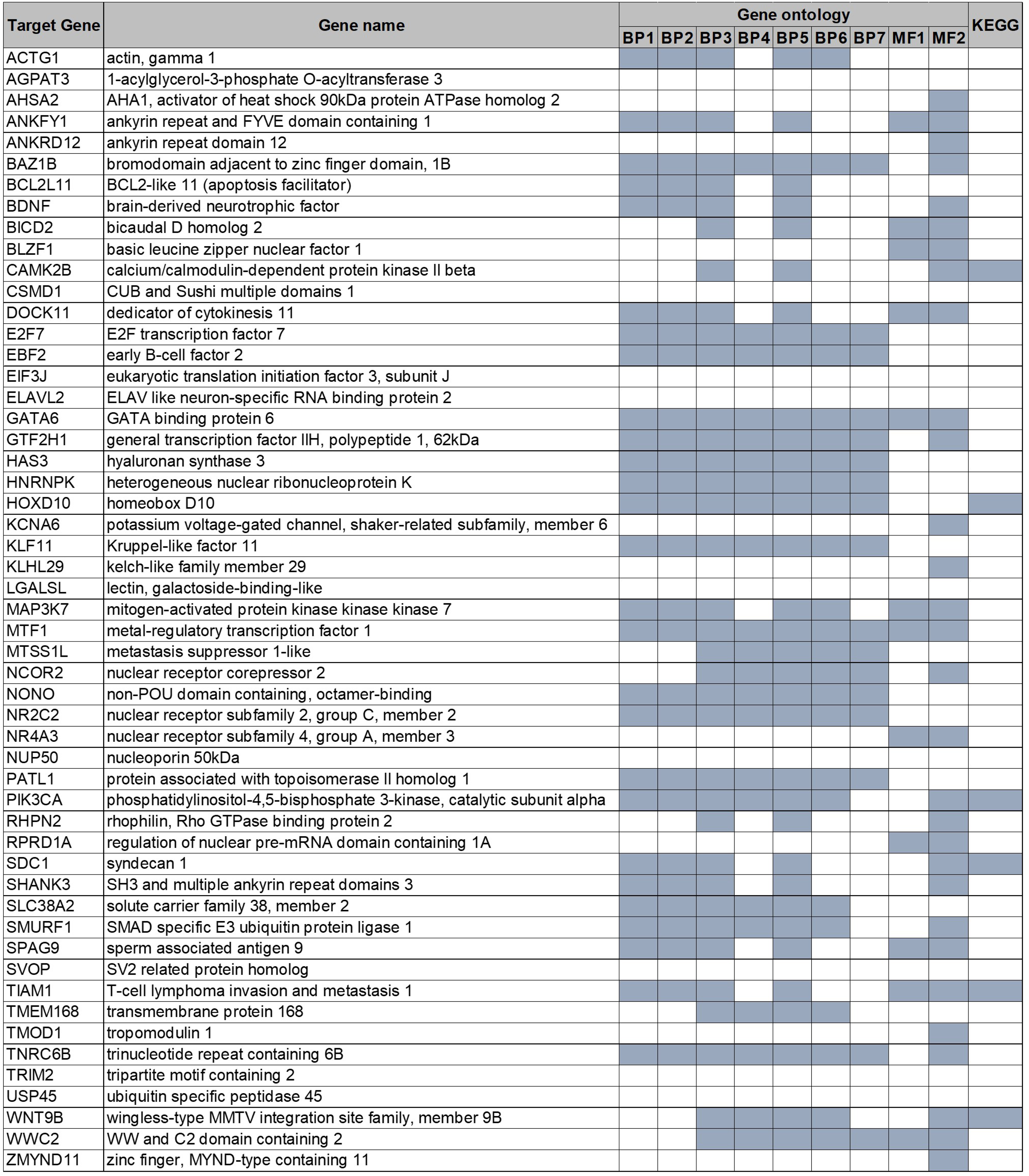

## Notes

### Competing Interest Statement

The authors have declared no competing interest.

